# Bacterial composition reflects fine-scale salinity changes while phylogenetic diversity exhibits a strong salt divide

**DOI:** 10.1101/2021.09.14.460410

**Authors:** Ariane L. Peralta, Mario E. Muscarella, Alexandra Stucy, Jo A. Werba, Michael W. McCoy

## Abstract

Climate change induced salinization events are predicted to intensify and lead to increased salt stress in freshwater aquatic ecosystems. As a consequence, formerly distinct abiotic conditions and associated biotic communities merge, and the emergence, loss, and persistence of microbial taxa modify the types and rates of ecosystem processes. This study examined how bacterial taxonomic and phylogenetic diversity and ecosystem function respond to acute salinization events where freshwater and estuarine communities and environments coalesce. We hypothesized that when the salinity change outpaces microbial adaptation or saline microbial populations are not yet established in formerly freshwater conditions, then these aquatic communities will exhibit diminished carbon cycling rates, decreased microbial diversity, and altered composition of microbial communities compared to historically freshwater communities. We used an experimental mesocosm approach to determine how salinity and the merging of distinct communities influenced resultant bacterial community structure and function. Each mesocosm represented different salinities (0, 5, 9, 13 psu). Two dispersal treatments, representing aquatic communities sourced from brackish 13 psu ponds and a mix of 13 psu and freshwater ponds, were added to all salinity levels and replicated four times. Results revealed that salinity, but not dispersal, decreased bacterial taxonomic and phylogenetic diversity. Carbon mineralization rates were highest in freshwater conditions and associated to bacterial taxa represented in low relative abundance. Acute salinity changes, such as localized flooding due to storm surge, will more negatively affect freshwater aquatic communities compared to chronic exposure to salinization where the communities have had time to adapt or turnover.

**IMPORTANCE STATEMENT:** Climate change induced salinization results in the mixing of formerly distinct environmental conditions and aquatic communities. This study examined the consequence of short-term, acute salinity stress on aquatic bacterial taxonomic and phylogenetic diversity and ecosystem function using an experimental approach. Results revealed that salinity, but not the source of aquatic communities, decreased bacterial taxonomic and phylogenetic diversity. Carbon mineralization rates, which represented ecosystem function, were highest in freshwater conditions and also associated with indicator bacterial taxa in low abundance relative to the total microbial community. Taken together, acute salinity changes will more negatively affect freshwater aquatic communities compared to chronic exposure to salinization where the communities have had time to adapt or turnover resulting in recovered biogeochemical functions.

## INTRODUCTION

Predicting microbial community response to environmental change depends on the tolerance and preference of microbial taxa to current and historical environmental conditions, ongoing changes in environmental conditions, and concurrent community assembly processes (1–3). Environmental changes, such as sea level rise and punctuated salinization events, are predicted to intensify in the coming decades as land development and climate change continue (4). Sealevel rise and saltwater intrusion in previously freshwater wetlands can diminish wetland ecosystem functions by increasing toxic sulfide production, decreasing inorganic nitrogen removal, and decreasing carbon storage. In some cases, diminished microbial functions negatively affect water quality, climate regulation, wetland accretion, and the health of wetland biota (5, 6). However, it is unclear how the composition and function of microbial communities at newly established fresh-salt water interfaces will change as salinization events increase in frequency and duration, especially with regard to subsets of taxa that respond to the coalescence event (i.e., the merging of distinct microbial communities and environments) (7, 8). As aquatic ecosystems shift from fresh to brackish waters, the concentration of alternative terminal electron acceptors (i.e., Fe(III), Mn(IV)) increases, which prompts CO_2_ production (6). As a result, environmental stressors are altering the microbial communities governing these carbon cycling rates.

Salinity tolerance has convergently evolved in unrelated lineages across the tree of life (9, 10). Salinity preference and osmotic tolerance are considered a complex microbial trait (11). Two mechanisms are responsible for osmotic or salinity stress tolerance in microorganisms. First, microorganisms can adapt by excluding harmful solutes (e.g., sodium, chloride) while gathering beneficial solutes necessary for metabolism by using active transport systems (12). Second, some microorganisms can create organic compounds to reduce the concentration gradient between the external environment and the cell resulting in the net export of unwanted solutes (13–15). Some studies suggest that long-term salinity changes, like those due to sea level rise, may occur slowly enough that the organisms can adapt and/or communities reassemble according to environmental changes. Long-term, incremental salinity increases have promoted microbial diversity and community establishment of salt-preference in a variety of taxa. Salinization in tidal and non-tidal systems results in different community responses. For example, tidal wetland decomposition rates increase as salinity increases due in part to marine/estuarine subsidies mixing with freshwater ecosystems (16). While experiencing short-term acute salinization, microbial populations have recovered from previously diminished carbon cycling functions (carbon dioxide and methane production) (17). Prior studies have revealed both increases in decomposition rates (15, 18) and decreases in decomposition rates (19, 20) as a result of increases in salinity. Previous studies on salinization effects of freshwaters reveal that punctuated salinity pulses to historically freshwater ecosystems, such as localized flooding due to storm surge in a climatic event, will more negatively affect freshwater communities compared to constant exposure to salinization over several years (2, 17, 21–24). The challenge to predicting salinization effects on decomposition rates is due to interactions among the following: direct effects of redox shifts between oxic and anoxic conditions, direct effects on heterotrophic communities and local nutrient availability, and the indirect effects due to changes in organic carbon sources (25).

Salinity tolerance and preference are traits that constrain microbial community response to salinization (26). As freshwater salinization across inland freshwaters persists, drastic changes to microbial community structure and function will continue to increase (27). The emergence, loss, and persistence of microbial taxa can alter the types and rates of ecosystem functions. The microbial taxa that will survive punctuated salinity changes will have specialized traits. These ‘effect’ traits can be directly linked to ecosystem functioning (11). Salinity preference is identified as one of the most complex trait measurements, classifying it based on how many genes are directly involved in coding the trait and how the trait is integrated with other mechanisms (11). Because specialized traits are needed for organisms to tolerate increasingly saline environments, major shifts in microbial composition and diversity are expected (28, 29). When freshwater and marine microbes are experimentally mixed, results showed that taxa which were rare at the initial inoculation became relatively abundant (7). As the community shifts, organisms that are better adapted to new conditions may become more active and abundant, which can alter the dominant ecosystem processes (1–3).

In this study, we address the research question: how does acute salinization affect microbial taxonomic and phylogenetic diversity and function when freshwater microbial communities mix with estuarine aquatic communities along freshwater to saltwater gradient? Specialized microbial traits are necessary to tolerate or thrive in saline environments. We hypothesize that during scenarios when salinity change outpaces microbial adaptation or saline microbial populations are not yet established in formerly freshwater conditions, aquatic communities will exhibit diminished carbon cycling rates (i.e., CO_2_ respiration) and decreased microbial diversity and altered the composition of microbial communities when compared to historically freshwater microbial communities. We also expect that microorganisms that persist in more saline conditions will be more phylogenetically related to each other (i.e., less genetic variation) compared to historically freshwater microorganisms. Dispersal of saline communities into freshwater ecosystems have the opportunity to establish and outcompete freshwater communities, but there is likely a period of transition where the mixing of environments and bacterial communities are dynamic and communities are reassembling. Understanding how salinization alters freshwater wetland bacterial phylogenetic-ecosystem function relationships can inform the management of carbon storage capacity in coastal wetlands experiencing increased salinization.

## MATERIALS AND METHODS

### Experimental Design

Microbial community composition and function were characterized based on aquatic samples that were collected from a replicated mesocosm experiment conducted from June-July 2015 (Figure 1) detailed in Werba et al. (30). To replicate a freshwater-saltwater aquatic system, peat moss (to serve as a nutrient pulse), sand (to serve as a benthic substrate), and water sourced from the municipal water supply were added to 150-gallon stock watering tanks (filled to 100 gallons). Each mesocosm pond was adjusted to one of 4 different salinities (measured in practical salinity units) using Instant Ocean® Sea Salt (4 levels: 0, 5, 9, 13 psu). Each tank was then seeded with an inoculation from natural ponds in the outer banks of North Carolina that matched the treatment’s psu (see **Supplemental Table S1** for coastal pond location and information). The inocula consisted of water, zooplankton and microbes. Twenty 1 L water samples were collected and filtered through 62.5 μm mesh across a one-hundred meter transect at each of the 5 coastal ponds and subsequently dispersed into each mesocosm tank. Bacterial communities represented in the source communities were likely particle- and zooplankton-associated because source communities were collected using a 62.5 μm mesh. Each tank was covered with a shade cloth to reduce the opportunity for other organisms from colonizing the mesocosms. Populations introduced via the initial inoculation were allowed to stabilize for 6 weeks before sampling began. After this acclimation period, mesocosms received additional colonists of zooplankton and microbes via simulated dispersal events from separate mesocosm ponds established as source tanks for salt (13 psu) and freshwater (0 psu) zooplankton and microbial communities. The source tanks served as dispersal treatments and consisted of either a saltwater (13 psu; salt dispersal treatment) community or a mix of communities from freshwater (0 psu) and saltwater (13 psu; fresh+salt dispersal treatment). Dispersal treatments consisted of 2 L of water from dispersal (source) tanks into the experiment (treatment) tanks. Sample collection occurred every 9 days (due to average time for completion of one zooplankton generation cycle) (31) over the 6-week mesocosm experiment for a total of six time points. The salinity × dispersal setup was replicated 4 times to generate 32 tanks (8 at each salinity level divided into 4 in each dispersal treatment) (30). For each tank, the following were measured using a YSI Pro: dissolved oxygen, ammonium (NH_4+_), temperature, and pH. Community analysis was focused on three of the time points (days 0, 18, and 45; 11 June 2015, 29 June 2015, and 25 July 2015, respectively), representing the initial, middle, and end of the experiment. Besides salinity levels, measured nitrogen and phosphorus concentrations were similar among treatment tanks (30).

**Figure 1.**
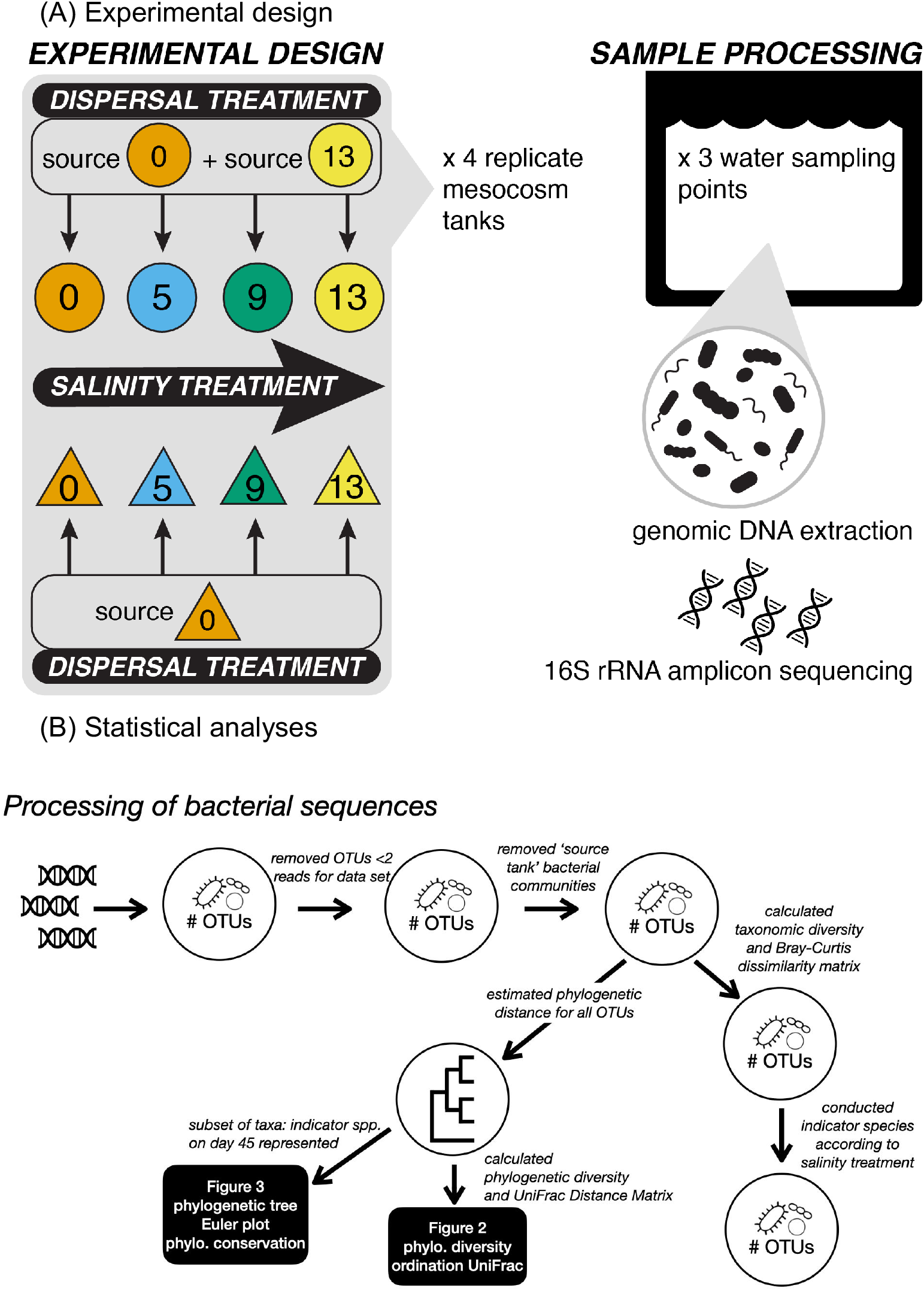
Schematic of experimental mesocosm to test the effect of salinity and dispersal of estuarine communities into freshwater coastal ponds (A) and bacterial sequence processing for statistical analyses (B).

### Microbial Sample Collection and Processing

Sample collection occurred every 9 days over the time of the 6-week mesocosm experiment for a total of six time points. During each sampling event, we collected 1 L of water from each tank. Each 1 L bottle of mesocosm water was homogenized and 200 mL of the water sample were concentrated onto 0.22 μm filters within 24 hours of field sampling (Supor-200; Pall Gelman, East Hills, NY) to collect microbial community samples. Filters were transferred into 2 mL sterile tubes and stored at -80° C until molecular analyses. We extracted DNA from filters collected at three (of the six) time points representing the initial, middle, final sampling dates (Days 0, 18, 45) using the PowerWater DNA Isolation Kit (MO BIO Laboratories).

### Microbial Community Sequencing

To examine shifts in microbial community composition and diversity, aquatic microbial communities in each mesocosm were characterized using paired-end targeted Illumina sequencing of the 16S rRNA gene (bacteria, archaea) (32). We extracted and purified the DNA found in the water of each mesocosm using the PowerWater DNA Isolation Kit (MO BIO Laboratories, Inc., Carlsbad, CA) collected on a 0.22 μm supor filter post-filtration. We used this DNA as a template in PCR reactions. To characterize bacterial communities, we used barcoded primers (515FB/806RB) originally developed by the Earth Microbiome Project (32) to target the V4 region of the bacterial 16S subunit of the ribosomal RNA gene (33, 34). PCR products were combined in equimolar concentrations and sequenced using paired-end (2×250 bp) approach using the Illumina MiSeq platform at Indiana University’s Center for Genomics and Bioinformatics.

Raw bacterial sequences were processed using the Mothur pipeline (version 1.41.3) (35). Briefly, paired-end contigs were assembled and quality trimmed, sequences were then aligned to the Silva Database (version 132) (36) chimeric sequences were removed using the VSEARCH algorithm (37), and we removed any sequences classified as Chloroplasts, Mitochondria, Archaea, or Eukaryotes. Next, we identified operational taxonomic units (OTUs) by separating sequences based on taxonomic class and then binning sequences using a 97% sequence similarity cutoff. Representative sequences from each OTU were used to generate a phylogenetic tree using FastTree v2.1 using the GTR model and CAT rate approximation (38). See Supplemental Material for full sequence processing pipeline.

### Compositional Analyses

We used a combination of taxonomic and phylogenetic approaches to characterize the diversity of bacterial communities in each mesocosm. First, we calculated alpha diversity (within-sample) using either Shannon’s diversity (taxonomic) or Faith’s Diversity (phylogenetic). The Faith’s Phylogenetic Diversity (PD) metric provides a simple measure of the phylogenetic relatedness of a community based on the summed branch lengths of its phylogenetic tree. We expected this measure to capture functional complementarity if more distantly related species are more functionally unique. We rarefied the OTU table to 20,000 observations before calculating alpha-diversity. Next, we calculated beta-diversity (among sites) by calculating Bray-Curtis distance for taxonomic diversity and weighted UniFrac distance for phylogenetic diversity. Differences in beta-diversity were visualized using Principal Coordinates Analysis.

### Carbon Mineralization Assay

On the final mesocosm sampling date, day 45, we measured the amount of CO_2_ respired from the aquatic communities using a laboratory-based bottle assay. Wheaton bottles (125 mL) fitted with septa were filled with water samples (25 mL) from each mesocosm tank. The CO_2_ concentration readings were determined using an LI-7000 Infrared Gas Analyzer (IRGA). On the first day (Day 0), bottles were filled with 25 mL of mesocosm tank water, and the gas samples were collected and analyzed immediately using the IRGA to determine the baseline CO_2_ concentration. A syringe was inserted into the septa and the headspace gas was mixed 3 times before pulling a sample and beginning analysis using the IRGA. This process was repeated on days one, three, and seven in order to determine CO_2_ respiration rates over time.

### Statistical Analyses

We completed statistical calculations in the R environment (R v4.1.2 R Core Development Team 2021). For bacterial diversity metrics, we conducted a linear mixed effects model with ‘date’, ‘salinity’, and ‘dispersal’ as fixed effects and ‘block’ as a random effect using the *lmer()* function in the lmerTest package (Kuznetsova et al. 2017) and the pbkrtest package (39). We ran linear mixed models that were fit using a restricted maximum likelihood (REML) approach and produced type II analysis of variances tables (ANOVA) tables based on the Kenward-Roger’s denominator degrees of freedom method using the *anova()* function. Then, we used principal coordinates analysis (PCoA) based on the Bray-Curtis dissimilarity of bacterial community composition and based on the weighted Unifrac distance of phylogenetic diversity along the salinity gradient. We ran a permutational multivariate analysis of variance (PERMANOVA) to examine among-treatment differences in bacterial communities. We identified which bacterial species were most representative of each salinity treatment using an indicator species analysis based on salinity level and also a categorical saline vs non-saline classification, and we included bacterial taxa with a relative abundance greater than 0.05 when summed across all plots. We performed PERMANOVA using the *adonis()* function in the vegan package (40) and the *indval()* function in the indicspecies package (41). We visualized overlap in indicator taxa across the salinity gradient using an Euler plot. To better understand the phylogenetic underpinnings of saline tolerant and sensitive taxa, we visualized the indicators on a phylogenetic tree and used the ConcenTrait algorithm to determine the phylogenetic depth at which saline tolerance/sensitivity is conserved (42). Briefly, this algorithm finds the nodes in the phylogeny at which 90% of the included tips (taxa) share the same trait. We implemented the ConcenTrait algorithm using a custom R script.

## Data Availability

All code and data used in this study are located in a public GitHub repository (https://github.com/PeraltaLab/CSI_Dispersal) and NCBI SRA BioProject ID PRJNA615001.

## RESULTS

We examined bacterial community turnover throughout the mesocosm experiment by comparing patterns in bacterial phylogenetic and taxonomic composition across the salinity gradient. Throughout the experiment, freshwater communities were phylogenetically distinct compared to saline communities based on weighted UniFrac distance metric (Fig. 2A, Table 1, PERMANOVA salinity: R^2^=0.172, P<0.001). Bacterial communities were phylogenetically similar (based on phylogenetic diversity) during day 18, but after 45 days at the experimental salinity level, communities at 9 and 13 psu were distinct from communities at 5 psu (Fig. 2C, Table 1, PERMANOVA salinity x date: R^2^=0.048, P<0.001). When bacterial taxonomic composition was compared, salinity was a much stronger environmental filter associated with bacterial community patterns based on taxonomy (Fig. 2B) compared to phylogeny (Fig. 2A)

**Figure 2.**
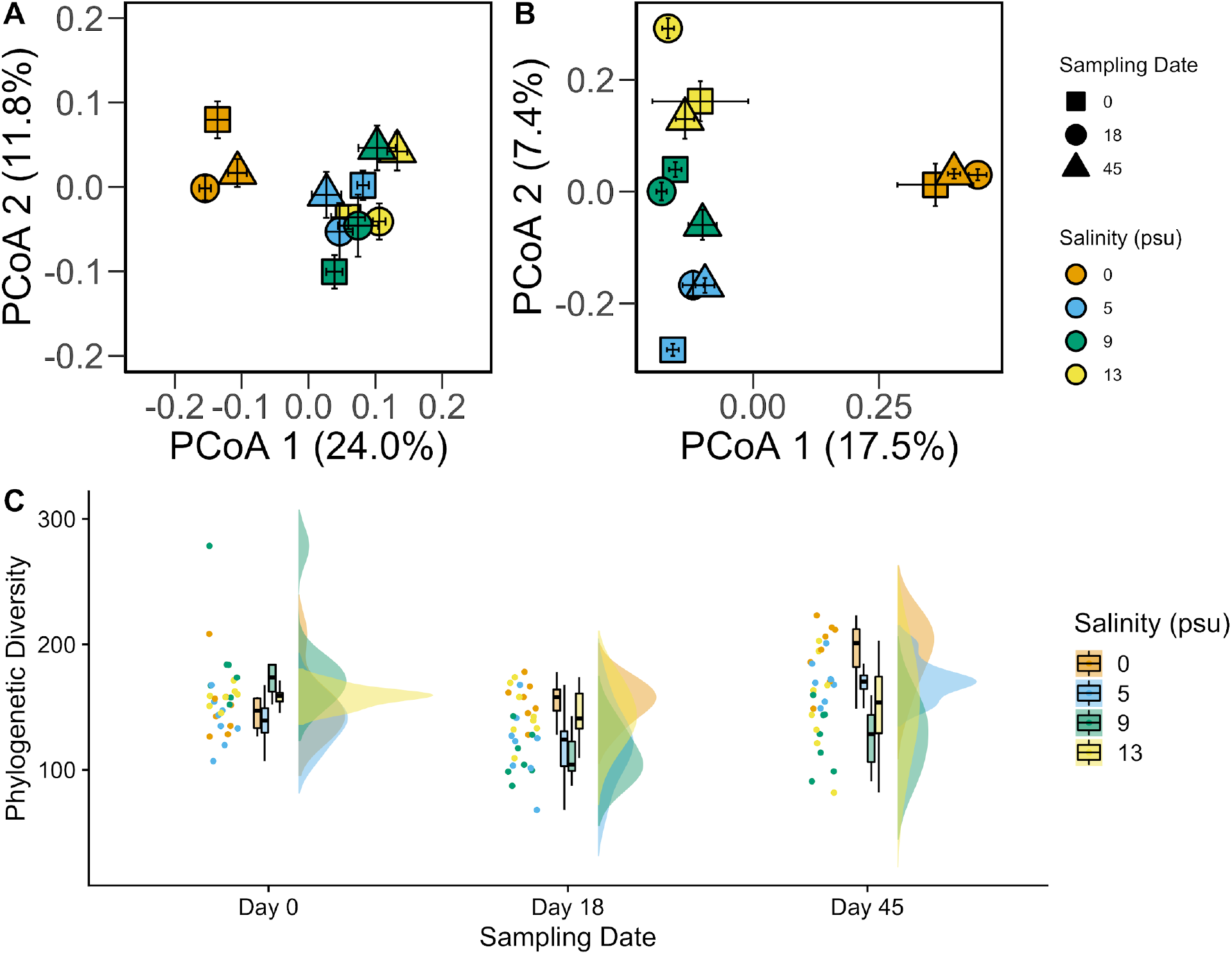
Bacterial phylogenetic and taxonomic patterns along a salinity gradient. Ordination based on a Principal Coordinates Analysis depicting bacterial phylogenetic distance (A) and Bray-Curtis dissimilarity distance (modified from Werba et al. 2020) (B), where symbols are colored according to salinity treatment (0, 5, 9, or 13 psu) and symbol shapes represent date of sampling (Day 0, 18, 45). Raincloud plots of phylogenetic diversity along a salinity gradient, where plots represent raw data depicted as filled circles (‘rain’), probability distributions (‘clouds’), and boxplots displaying mean and 95% confidence intervals (C). Colors represent salinity treatment (orange=0 psu, blue=5 psu, green=9 psu, yellow=13 psu), and shapes represent date of sampling (square=Day 0, circle=18, triangle=45).

**Table 1.** Summary PERMANOVA output comparing bacterial communities due to the factors salinity, dispersal, and sampling date.

| Factor | Df | Sums Sq | Mean Sq | F.Model | R<sup>2</sup> | Pr(>F) |
| --- | --- | --- | --- | --- | --- | --- |
| Salinity | 1 | 1.275 | 1.275 | 28.576 | 0.172 | **<0.001** |
| Dispersal | 4 | 0.210 | 0.052 | 1.175 | 0.028 | 0.200 |
| Date | 2 | 0.515 | 0.258 | 5.772 | 0.069 | **<0.001** |
| Salinity:Dispersal | 1 | 0.041 | 0.041 | 0.916 | 0.006 | 0.509 |
| Salinity:Date | 2 | 0.353 | 0.177 | 3.956 | 0.048 | **<0.001** |
| Dispersal:Date | 8 | 0.290 | 0.036 | 0.811 | 0.039 | 0.914 |
| Salinity:Dispersal:Date | 2 | 0.063 | 0.032 | 0.707 | 0.008 | 0.858 |
| Residuals | 105 | 4.686 | 0.045 | 0.6304 |  |  |
| Total | 125 | 7.433 | 1 |  |  |  |

Next, we compared patterns in bacterial phylogenetic diversity to taxonomic composition across the salinity gradient. Communities observed in the freshwater conditions increased in mean PD over the 45 day experiment. The PD of communities at the transition 9 psu tanks declined throughout the experiment duration, while PD of communities in the 13 psu tanks increased in variability from day 0 to 45 (Fig. 2C, Table 2, ANOVA, F_2,79_=5.997, P=0.004). After 45 days, PD was highest in freshwater and lowest in brackish tanks (Fig. 2C).

**Table 2.** Summary of Type II Analysis of Variance Table with Kenward-Roger’s method comparing phylogenetic diversity. We conducted a linear mixed effects model with ‘date’, ‘salinity’, and ‘dispersal’ as fixed effects and ‘block’ as a random effect.

| Factor | Df | Sum Sq | Mean Sq | F-value | Pr(>F) |
| --- | --- | --- | --- | --- | --- |
| Salinity | 1 | 3505 | 3505.3 | 3.703 | **0.058** |
| Dispersal | 1 | 79 | 79.4 | 0.084 | 0.773 |
| Date | 2 | 14396 | 7197.9 | 7.604 | **<0.001** |
| Dispersal:Salinity | 1 | 265 | 264.6 | 0.280 | 0.598 |
| Dispersal:Date | 2 | 901 | 450.4 | 0.476 | 0.623 |
| Salinity:Date | 2 | 11353 | 5676.5 | 5.997 | **0.004** |
| Dispersal:Salinity:Date | 2 | 136 | 67.8 | 0.072 | 0.931 |
| Residuals | 79 | 74778 | 946.6 |  |  |

There are OTUs represented across major phyla that are associated with all salinity levels (Fig. 3A). Based on the unrooted tree (Fig. 3A), we observed that the smallest number of OTUs were represented at 9 psu treatment compared to other treatments. Next, we used a series of Euler plots to reveal shared and unique indicator OTUs. A list of “indicators” that were significantly associated with each salinity treatment is found in Table S2. These indicators, reported at the class-level, are the taxonomic groups best associated with each treatment. Based on OTUs identified as indicator taxa, 254 OTUs are unique to the freshwater treatment, 135 OTUs are unique to the 5 psu treatment, 35 OTUs were unique to the 9 psu treatment, and 124 OTUs are unique to the 13 psu treatment (Fig. 3B). When the OTUs were grouped into salt or no salt, there are 206 observed indicator OTUs shared across 5, 9, or 13 psu environments, and 257 OTUs unique to the 0 psu treatment (Fig. 3B). Finally, when examining the depth of phylogenetic relatedness, results from the ConcenTrait analysis revealed that saline vs non-saline microbes exhibit a similar depth in average phylogenetic relatedness (Fig. 3C).

**Figure 3.**
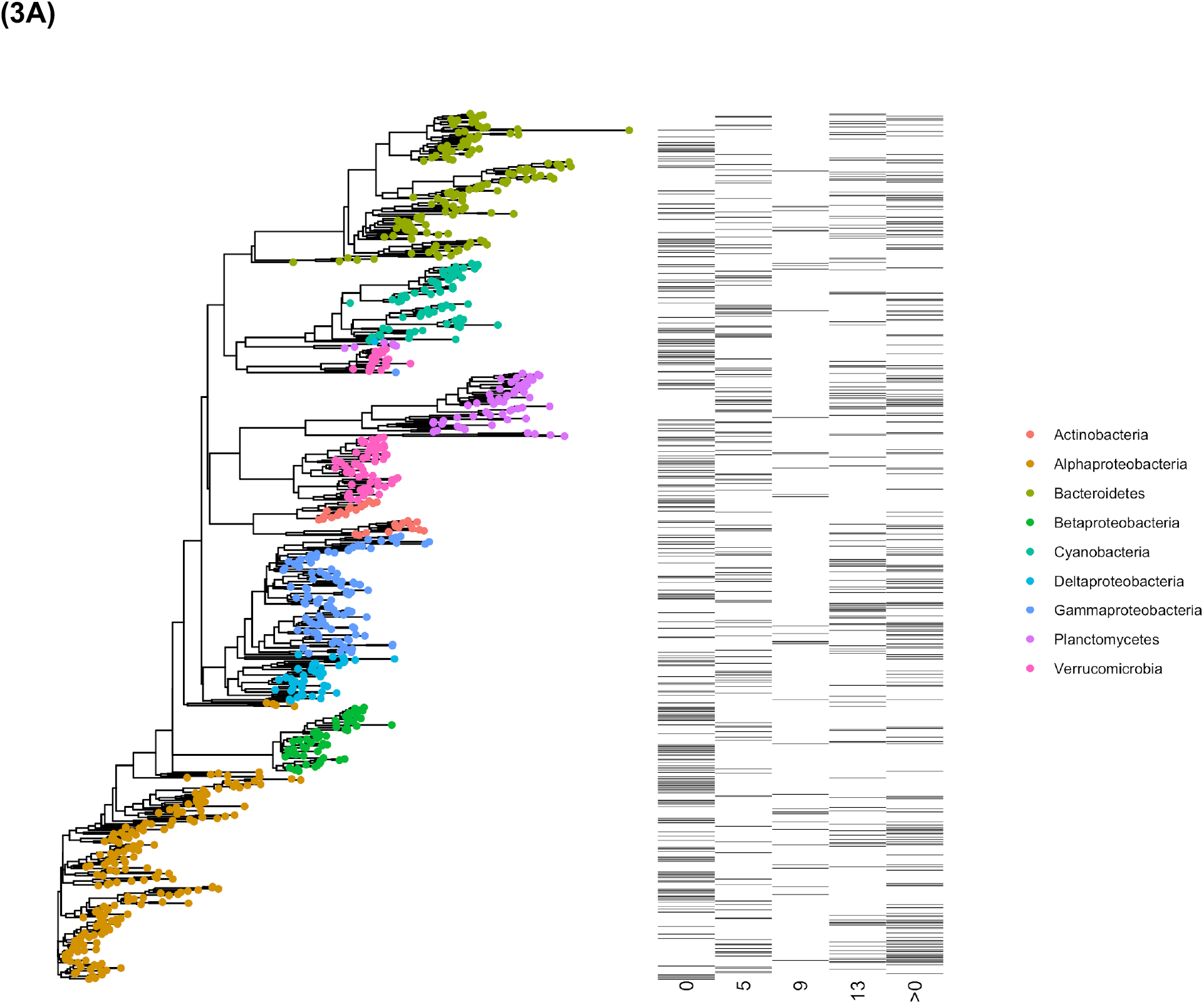
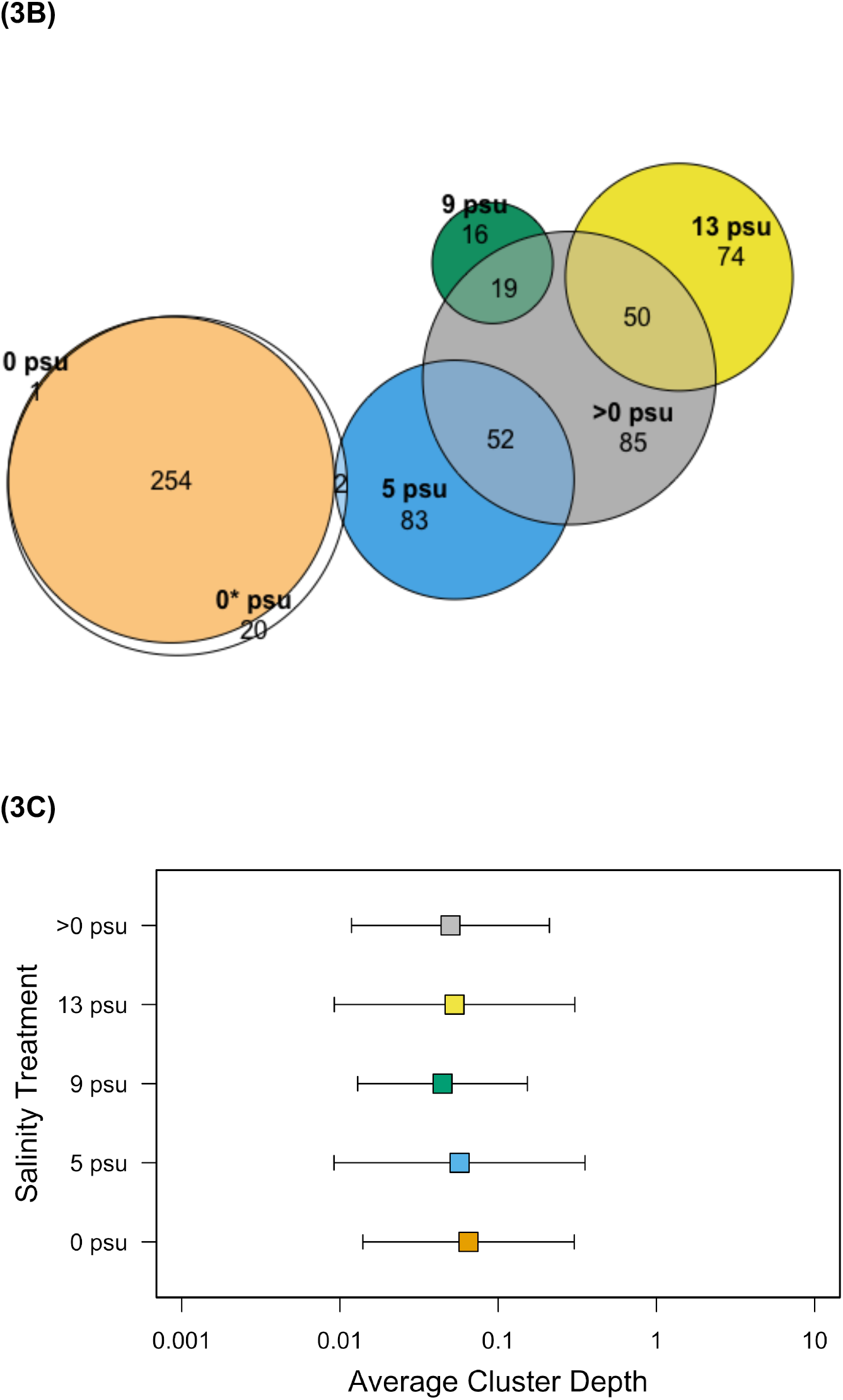
Phylogenetic patterns of indicator bacterial taxa. Phylogenetic tree of bacterial taxa that were identified to be statistically associated with salinity treatment (0, 5, 9, 13 psu or combined salt treatment represented by >0 psu); each taxon tip color corresponds to phyla level grouping and is associated with a heat map which represents OTU relative abundance salinity treatment (A). Euler plots represent bacterial OTU that are unique or shared across salinity treatments. The 0* label refers to combined salt treatments (combining 5, 9, 13 psu into a single ‘salt’ treatment).(A). Average cluster depth estimated using ConcenTrait used to depict phylogenetic distance of bacterial assemblages observed across the salinity gradient (C).

We examined the relationship between carbon mineralization and phylogenetic diversity and relative abundance (Fig. 4A). At the transition 9 psu treatments, we measured the lowest carbon mineralization values compared to other salinity treatments. Carbon mineralization rates were highest in freshwater conditions and associated with indicator bacterial taxa that were measured at low relative abundance (< 0.01%) (Fig. 4B). In contrast, the relative abundance of indicator taxa represented in the saltwater conditions represented > 25% of the community.

**Figure 4.**
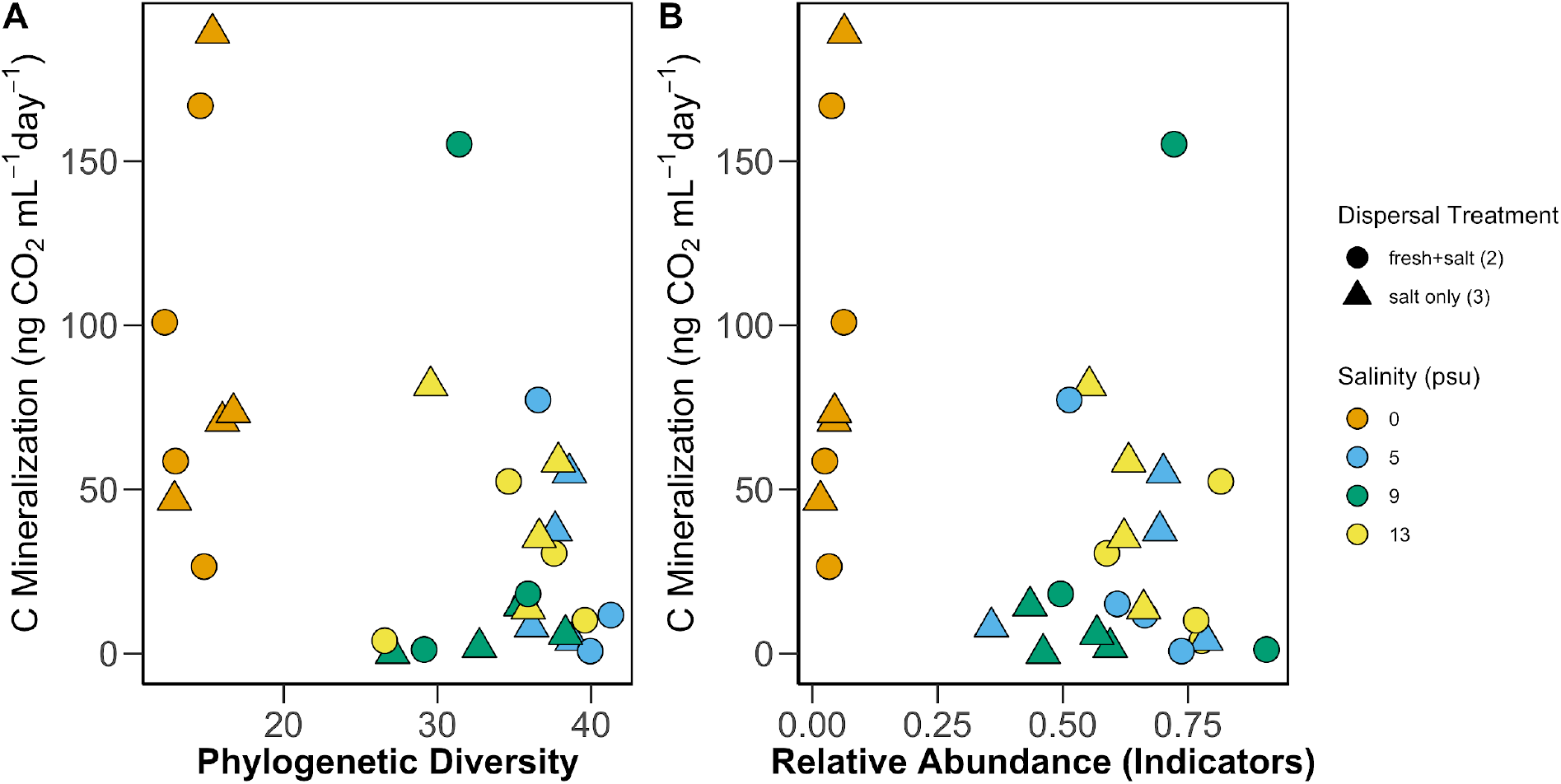
Relationship between carbon mineralization and bacterial phylogenetic diversity (PD), and taxonomic-based community metrics (relative abundance) for Day 45 (last day of experiment).

## DISCUSSION

This study specifically focused on determining the extent that short-term salinization events influenced reassembly patterns of aquatic communities and carbon mineralization rates. Results revealed that salinity strongly influenced bacterial phylogenetic diversity; bacterial communities tended to respond to salinity, rather than dispersal, because it is such a strong environmental filter (7, 10). Past research has shown that strong environmental filters produce phylogenetically clustered communities (11, 43, 44). However, the mixing of two formerly distinct communities and their environments could affect community structure and function in unpredictable ways. Outcomes depend on the strength of the environmental change and the capacity for taxa to successfully establish and reproduce after mixing events occur (2, 3). There are many factors that determine how bacterial community composition will change in response to an influx of saline-dominant communities following a salinization event. Both stochastic processes (such as drift) and deterministic processes (such as the interaction of biotic and abiotic factors) within an ecosystem can determine patterns of community assembly via four processes: selection, dispersal, diversification, and drift (43, 45). Abiotic factors in an environment often act as environmental filters, selecting for organisms that are adapted to establish and persist. In this study, once mesocosms were seeded with aquatic inocula, the environmental conditions strongly dictated the newly established community assemblages based on taxonomy (Fig. 2B and (30)) but not phylogeny (Fig. 2A).

Bacterial taxa were more related to each other in transition salinity environments (5 and 9 psu) compared to fresh and the most saline endpoint environments. The composition of the bacterial community (based on phylogeny) was similar in the 5 and 9 psu treatment, but the phylogenetic diversity rebounded for communities in the 5 psu compared to 9 psu treatments. The 9 psu environment could be particularly stressful for the resorted ‘transition’ community. In the current study, we added a mixture of freshwater and mesohaline (13 psu) aquatic communities in equal proportion as dispersal treatments along an experimental salinity gradient. The dispersal treatments represented natural dispersal methods of aquatic communities via water; a fresh community being mixed with a saline community can occur in storm events with coastal flooding, as well as gradual mixing of these environments via sea level rise. These methods of dispersal are on different time-scales, but each is potentially important to community assembly at mixing zones. Because dispersal of microbes is often a passive process, we recognize that the longest transport of microbial species is via wind, water, and attachment to mobile organisms (43).

Overall, bacterial communities tended to respond to salinity, rather than dispersal. Our results provide further support that salinity is a strong environmental filter especially in light of salinity tolerance being a relatively deeply conserved trait (10, 11, 46). The depth of phylogenetic relatedness was similar for saline and non-saline communities. This suggests that there is a deep fresh-salt divide as previously revealed (7, 8, 10).

Since salinity tolerance is considered a phylogenetically conserved trait, it is more likely that the reassembly of the mixed community to represent a new state. Past studies show that selection may act on traits that are subject to HGT, which can alter processes by transferring genetic material (i.e., a trait that would allow a previously rare organism to establish and persist in a previously inhospitable environment) (47, 48). These organisms can now explore new fitness landscapes and diversify (43). However, gaining a trait is not always a simple process, as some traits are complex and cannot be easily inherited (10, 11, 46). This study did not support the idea that selection for salinity tolerance was occurring due to the short-term study design. Ecological drift (constant changes in relative abundance of organisms) can potentially affect community assembly. In particular, low abundance organisms (i.e., majority of microbial species) are more vulnerable to drift (43). Alternatively, rare taxa that were not detectable based on amplicon sequencing methods were identified as responding positively to mixing events, making the role of rare taxa important for driving community composition changes (7, 8).

Salinity stressors on freshwater aquatic communities could indirectly influence bacterial communities through influence other members of the food web. For example, microeukaryotes such as protists could shift in composition, changing grazer pressures on bacterial prey (49, 50). While the current study did not examine the other members of the food web, past studies revealed the importance of eco-evolutionary processes of food web interactions that have important implications for climate change effects on carbon cycling (51, 52).

It is also important to consider the intensity and length of the disturbance affecting a microbial community (53). The disturbance in this experiment was the addition of salinity and saline-adapted aquatic species to historically freshwater communities. This disturbance was intended to represent salinization due to acute events such as storm surge. At the end of the experiment, phylogenetic diversity (PD) was low for both dispersal treatments at the highest salinity, indicating that the bacteria persisting in those tanks were more related to each other (lower PD). This observation supported the hypothesis that PD would be lowest in the most saline tank, due to salinity tolerance being a complex trait.

After saline and freshwater communities mixed, we observed the highest carbon mineralization rates in freshwater environments and associated disproportionately lower abundance taxa. In contrast, microbes carried out lower carbon mineralization rates under saline conditions, which were associated with indicator taxa at >25% relative abundance. Past studies showed that conditionally rare microbial taxa play disproportionate role in community patterns over time and considered keystone taxa important for ecological responses to disturbance (9– 11). Salinization influences microbes through osmotic stress and ion-specific toxicity, but salinization can also provide terminal electron acceptors to drive and change the rates and types of microbial metabolisms. Some studies suggest that long-term salinity changes, like those due to sea level rise, may occur slowly enough that the organisms can adapt (e.g., (2)). Acute salinity changes caused by “pulses” of salinity influx, such as localized flooding due to storm surge, will more negatively affect freshwater aquatic communities compared to chronic exposure to salinization where the communities have had time to adapt or turnover resulting in recovered biogeochemical functions (e.g., (2)). Thus, unexpected pulses of salinity are observed (in some cases) to cause greater damage to ecosystem structure and function than long-term salinity changes due to the inability of organisms to adapt or change quickly following one of these events (6, 17, 54). The current study provides insight into salinization effects of saline communities mixing with naïve freshwater communities, an occurrence that is expected at the early onset of salinization events (e.g., sea-level rise, intense storm surges upriver).

Global climate change plays a large role in facilitating increased dispersal of microbes; the predicted increases in extreme weather events may potentially promote increased spread of microbes to new, adaptable environments (55, 56). As previously discussed, microbes have several mechanisms for adapting to environmental change; this experiment did not address which mechanisms were used. Future studies focused on bacterial community structure and function under increasing salinity would benefit from examining mechanisms of adaptation that taxa may use when placed under environmental stress.

## Acknowledgments

We thank Spencer Wilkinson, Amanda Dunn, and Mary-Grace Lee for help with data collection. We also thank Mike Piehler and Corey Adams for logistical help at the Coastal Studies Institute. This work was supported in part by the East Carolina University Division of Research, Economic Development, and Engagement and the Department of Biology.

## Notes

https://github.com/PeraltaLab/CSI_Dispersal

### Competing Interest Statement

The authors have declared no competing interest.

### Summary of Updates

We clarified experimental design text and results/discussion highlights for clarity in this version of the manuscript. Tables 1 and 2 were moved from supplemental into main paper. Supplemental files updated.

