## Supplemental Tables for "Bacterial composition reflects fine-scale salinity changes while phylogenetic diversity exhibits a strong salt divide"

[1]Department of Biology, East Carolina University, Howell Science Complex, Greenville, NC 27858; [2]Department of Biology and Wildlife, Institute of Arctic Biology, University of Alaska Fairbanks, Fairbanks, AK, 99775; [3]Department of Biology, McMaster University, 1280 Main Street West, Hamilton, Ontario, Canada, L8S 4K1; [4]Harbor Branch Oceanographic Institute, Florida Atlantic University, Fort Pierce, FL, 34946, USA

*corresponding author

**Supplemental Materials | TABLES**

**Supplemental Table S1.** Site characteristics of the coastal ponds used as sources for freshwater and saltwater dispersal sources. Table displays the site name, date of collection, GPS location, temperature in degrees Celsius, salinity on date of collection, and pH on date of collection.

| Site | Collection Date | GPS Location | Temperature (°C) | Salinity | pH |
|---|---|---|---|---|---|
| CSI 1 | 3 May 2015 | N 35°52.443', W 075°39.640 | 29.7 | 0.20 | 9.65 |
| A3 | 3 May 2015 | N 35°36.632' W 075°28.074' | 30.2 | 5.20 | 9.29 |
| Bodie 1 | 3 May 2015 | N 35°49.207' W 075°33.755' | 31.9 | 9.46 | 8.59 |
| A2 | 3 May 2015 | N 35°41.89' W 075°29.041' | 28.4 | 15.12 | 7.94 |

**Supplemental Table S2.** Indicator operational taxonomic unit (OTU at class-level) for each salinity treatment. Values in parentheses indicate how many indicator OTUs were associated with each class.

| 0 psu | 5 psu | 9 psu | 13 psu |
|---|---|---|---|
| Actinobacteria (10) | Actinobacteria (5) | Alphaproteobacteria (14) | Actinobacteria (3) |
| Alphaproteobacteria (69) | Alphaproteobacteria (25) | Betaproteobacteria (1) | Alphaproteobacteria (30) |
| Betaproteobacteria (30) | Betaproteobacteria (7) | Cyanobacteria (3) | Betaproteobacteria (4) |
| Cyanobacteria (18( | Cyanobacteria (21) | Cytophagia (6) | Cyanobacteria (6) |
| Cyanobacteria/Chloroplast_uncl (2) | Cytophagia (8) | Deltaproteobacteria (1) | Cytophagia (8) |
| Cytophagia (15) | Deltaproteobacteria (11) | Flavobacteriia (1) | Deltaproteobacteria (3) |
| Deltaproteobacteria (15) | Flavobacteriia (5) | Gammaproteobacteria (5) | Flavobacteriia (7) |
| Flavobacteriia (5) | Gammaproteobacteria (18) | Planctomycetia (1) | Gammaproteobacteria (32) |
| Gammaproteobacteria (24) | Opitutae (3) | Spartobacteria (1) | Opitutae (3) |
| Opitutae (12) | Phycisphaerae (2) | Verrucomicrobiae (3) | Phycisphaerae (2) |
| Planctomycetia (14) | Planctomycetia (16) | | Planctomycetia (13) |
| Spartobacteria (4) | Sphingobacteriia (7) | | Sphingobacteriia (12) |
| Sphingobacteriia (13) | Subdivision3 (4) | | Verrucomicrobiae (2) |
| Subdivision3 (3) | Verrucomicrobiae (6) | | |
| Verrucomicrobia_uncl (4) | | | |
| Verrucomicrobiae (15) | | | |

**Supplemental Materials | Sequence Processing Pipeline**

All sequence processing was done using mothur (v. 1.41.3; mothur citation) and the following pipeline:

1. Raw reads were assembled into contigs and screened based on length and ambiguous base calls.

```
make.contigs(file=CSI.files, processors=12, inputdir=../raw/,
outputdir=../analysis/)
screen.seqs(fasta=CSI.trim.contigs.fasta, group=CSI.contigs.groups,
summary=CSI.trim.contigs.summary, maxambig=0, maxlength=275)
unique.seqs(fasta=CSI.trim.contigs.good.fasta)
```

2. Sequences were aligned to the Silva reference database (v. 128)

```
count.seqs(name=CSI.trim.contigs.good.names,
group=CSI.contigs.good.groups)
align.seqs(fasta=CSI.trim.contigs.good.unique.fasta,
reference=~/dbs/silva.v4.fasta)
summary.seqs(fasta=CSI.trim.contigs.good.unique.align,
count=CSI.trim.contigs.good.count_table)
```

3. The aligned sequences were then screened based on alignment quality and trimmed

```
screen.seqs(fasta=CSI.trim.contigs.good.unique.align,
count=CSI.trim.contigs.good.count_table,
summary=CSI.trim.contigs.good.unique.summary, start=1968, end=11550,
maxhomop=8)
```

```
summary.seqs(fasta=CSI.trim.contigs.good.unique.good.align,
count=CSI.trim.contigs.good.good.count_table)
filter.seqs(fasta=CSI.trim.contigs.good.unique.good.align, vertical=T, trump=.)
unique.seqs(fasta=CSI.trim.contigs.good.unique.good.filter.fasta,
count=CSI.trim.contigs.good.good.count_table)
```

4. Sequences were then pre-clustered using a 2 bp differences criteria

```
pre.cluster(fasta=CSI.trim.contigs.good.unique.good.filter.unique.fasta,
count=CSI.trim.contigs.good.unique.good.filter.count_table, diffs=2)
summary.seqs(fasta=CSI.trim.contigs.good.unique.good.filter.unique.precluster.f
asta,
count=CSI.trim.contigs.good.unique.good.filter.unique.precluster.count_table)
```

5. Chimeric sequences were removed using the vsearch algorithm

```
chimera.vsearch(fasta=CSI.trim.contigs.good.unique.good.filter.unique.precluster
.fasta,
count=CSI.trim.contigs.good.unique.good.filter.unique.precluster.count_table,
dereplicate=t)
remove.seqs(fasta=CSI.trim.contigs.good.unique.good.filter.unique.precluster.fas
ta, count=CSI.trim.contigs.good.unique.good.filter.unique.precluster.count_table,
accnos=CSI.trim.contigs.good.unique.good.filter.unique.precluster.denovo.vsear
ch.accnos)
```

6. Sequences were classified using the RDP database and undesired lineages were removed

```
classify.seqs(fasta=CSI.trim.contigs.good.unique.good.filter.unique.precluster.pic
k.fasta,
count=CSI.trim.contigs.good.unique.good.filter.unique.precluster.pick.count_tabl
e, reference=~/dbs/trainset16_022016.pds.fasta,
taxonomy=~/dbs/trainset16_022016.pds.tax, cutoff=80)
remove.lineage(fasta=CSI.trim.contigs.good.unique.good.filter.unique.precluster.
pick.fasta,
count=CSI.trim.contigs.good.unique.good.filter.unique.precluster.pick.count_tabl
e,
taxonomy=CSI.trim.contigs.good.unique.good.filter.unique.precluster.pick.pds.wa
ng.taxonomy, taxon=Archaea-Mitochondria-
Cyanobacteria/Chloroplast;Chloroplast;-unknown-Eukaryota)
```

7. Sequences were then clustered. First, sequences were split using high level taxonomic identification and then the optiClust algorithm was used to cluster at 97% sequence similarity.

```
cluster.split(fasta=CSI.trim.contigs.good.unique.good.filter.unique.precluster.pick.
pick.fasta,
count=CSI.trim.contigs.good.unique.good.filter.unique.precluster.pick.pick.count_
table,
taxonomy=CSI.trim.contigs.good.unique.good.filter.unique.precluster.pick.pds.wa
ng.pick.taxonomy, splitmethod=classify, taxlevel=4, cutoff=0.03)
```

8. Last, a taxa-by-site matrix was created and representatives were picked for each OTU

```
make.shared(list=CSI.trim.contigs.good.unique.good.filter.unique.precluster.pick.
pick.opti_mcc.list,
count=CSI.trim.contigs.good.unique.good.filter.unique.precluster.pick.pick.count_
table, label=0.03)
classify.otu(list=CSI.trim.contigs.good.unique.good.filter.unique.precluster.pick.pi
ck.opti_mcc.list,
count=CSI.trim.contigs.good.unique.good.filter.unique.precluster.pick.pick.count_
table,
taxonomy=CSI.trim.contigs.good.unique.good.filter.unique.precluster.pick.pds.wa
ng.pick.taxonomy, label=0.03)
get.oturep(list=CSI.trim.contigs.good.unique.good.filter.unique.precluster.pick.pic
k.opti_mcc.list,
fasta=CSI.trim.contigs.good.unique.good.filter.unique.precluster.pick.pick.fasta,
count=CSI.trim.contigs.good.unique.good.filter.unique.precluster.pick.pick.count_
table, method=abundance, label=0.03)
```
